# Heuristic Spike Sorting Tuner (HSST), a framework to determine optimal parameter selection for a generic spike sorting algorithm

**DOI:** 10.1101/2020.05.21.108902

**Authors:** David A. Bjånes, Lee E. Fisher, Robert A. Gaunt, Douglas J. Weber

**Affiliations:** Rehab Neural Engineering Labs, University of Pittsburgh, Pittsburgh, USA; Department of Electrical and Computer Engineering, University of Pittsburgh, Pittsburgh, USA; Department of Physical Medicine and Rehabilitation, University of Pittsburgh, Pittsburgh, USA; Department of Bioengineering, University of Pittsburgh, Pittsburgh, USA; Center for the Neural Basis of Cognition, the University of Pittsburgh Brain Institute, University of Pittsburgh, Pittsburgh, USA; McGowan Institute for Regenerative Medicine, University of Pittsburgh, Pittsburgh, USA

**Keywords:** neurophysiology, spike sorting, heuristics, metrics, single-unit

## Abstract

Heuristic Spike Sorting Tuner (HSST), a framework to determine optimal parameter selection for a generic spike sorting algorithm. *bioRxiv* First published May 21, 2020. Extracellular microelectrodes frequently record neural activity from more than one neuron in the vicinity of the electrode. The process of labeling each recorded spike waveform with the identity of its source neuron is called spike sorting and is often approached from an abstracted statistical perspective. However, these approaches do not consider neurophysiological realities and may ignore important features that could improve the accuracy of these methods. Further, standard algorithms typically require selection of at least one free parameter, which can have significant effects on the quality of the output. We describe a Heuristic Spike Sorting Tuner (HSST) that determines the optimal choice of the free parameters for a given spike sorting algorithm based on the neurophysiological qualification of unit isolation and signal discrimination. A set of heuristic metrics are used to score the output of a spike sorting algorithm over a range of free parameters resulting in optimal sorting quality. We demonstrate that these metrics can be used to tune parameters in several spike sorting algorithms. The HSST algorithm shows robustness to variations in signal to noise ratio, number and relative size of units per channel. Moreover, the HSST algorithm is computationally efficient, operates unsupervised, and is parallelizable for batch processing.

**NEW & NOTEWORTHY:** HSST incorporates known neurophysiological priors of extracellular neural recordings while simultaneously taking advantage of powerful abstract mathematical tools. Rather than simply selecting free parameters prior to running a sorting algorithm, HSST executes a sorting algorithm across a range of input parameters, using heuristic metrics to detect which spike-sorting output is most physiologically plausible. This novel approach enables unsupervised spike-sorting exceeding the performance of previous methods, thereby enabling the processing of large data sets with confidence.

## INTRODUCTION

Investigations in the central and peripheral nervous systems often focus on extracting information about the spiking activity of individual neurons. When extracellular electrodes are used to record this activity, action potentials, or spikes, can be recorded simultaneously from multiple neurons in the immediate vicinity of a single electrode tip (Lemon, 1984; Pedreira, Martinez, Ison, & Quian Quiroga, 2012). Importantly, neurons recorded from the same electrode may exhibit very different patterns of activity and be correlated to different behavioral features. Each neuron produces a stereotypical voltage waveform at the electrode based on physical factors including cell geometry, neuron-electrode distance and tissue impedance (Camuñas-Mesa & Quiroga, 2013; Gold, 2006; Hild & Tasaki, 1962). Determining which waveforms are associated with the same source neuron is a process known as spike sorting (Chen, Carlson, & Carin, 2011; Gerstein & Clark, 1964; Hill, Mehta, & Kleinfeld, 2011; Keehn, 1966; Letelier & Weber, 2000; Lewicki, 1999; Prochazka, Conrad, & Sindermann, 1972; Quiroga, Nadasdy, & Ben-Shaul, 2004; Shoham, Fellows, & Normann, 2003; Yuan, Yang, & Si, 2012). Only after the waveforms are sorted can hypotheses be tested that depend on the temporal behavior of those neurons (Hubel & Wiesel, 1959).

Spike sorting is frequently approached as a multi-step process and is broken down into approximately three steps: 1) spike detection, 2) feature extraction, and 3) clustering (Gibson, Judy, & Markovic, 2008). Many techniques and algorithms have been developed to address the spike sorting problem (Abeles & Goldstein, 1977; Lewicki, 1999; Rey, Pedreira, & Quian Quiroga, 2015). These methods are typically supervised (i.e. manual sorting by visual waveform inspection), semi-supervised (i.e. manual inspection of automatically generated sorts) or unsupervised (i.e. sorting performed purely algorithmically). All approaches have specific strengths and weaknesses. Manual sorting is typically based on visual inspection, which can lead to inter- and intra-sorter variability (Harris, Henze, Csicsvari, Hirase, & Buzsáki, 2000; Wood, Black, Vargas-Irwin, Fellows, & Donoghue, 2004). Manual sorting is also extremely time intensive and impractical for applications with large datasets. Supervised or semi-supervised methods avoid these pitfalls, but often fail when sorting spikes that have a low signal-to-noise ratio (SNR). Nevertheless, manual or supervised techniques remain common practice in many labs.

Sorting algorithms necessarily have parameters that must either be set explicitly by the user or automatically by the algorithm itself. These parameters are often selected to reflect underlying assumptions about the data (e.g. expected number of neurons, variance in waveform shapes). For unsupervised algorithms, correctly estimating these parameters largely determines performance; poorly chosen parameters can create erroneous results, with waveforms from the same neuron classified into separate groups (over-sorting) or waveforms from multiple neurons classified together (under-sorting). Many common sorting algorithms suffer from these problems, requiring data to be inspected manually and resorted, effectively losing the advantage of an unsupervised method.

For a truly unsupervised method to sort a wide variety of data, the algorithm must accurately estimate these parameters, adding to the complexity of classification. Often the choice of parameters results in either an unrealistically large numbers of clusters with low waveform variability or a small number of clusters with unreasonably high waveform variability. It is difficult or impossible to determine a priori how many true neurons are present in a recording, since many of the measurable features (including the SNR, relative size of the proposed clusters, variance of waveforms within a cluster, etc.) are highly variably across datasets. For a basic example, the K-Means algorithm (MacQueen, 1967) has a single parameter *K*, which fixes the number of clusters discovered by the algorithm. Setting an incorrect value for *K* results in over- or under-sorting errors. To solve this problem, many algorithms utilize purely mathematical techniques to optimize the parameter selection process. By fixing constraints on the relationship between waveform variability and the number of clusters, these algorithms generally use statistical optimization methods to fit the parameters. Many use dimensionality reduction techniques to optimize parameter selection (Hulata, Segev, & Benjacob, 2002). To separate the waveforms, they collapse data variability onto a lower dimensional space. However, the underlying neural dynamics which govern spike generation are often lost or disregarded through this process. Rather than discard this information, we seek to utilize it by building a series of heuristic metrics to judge the sort quality of the output of a sorting algorithm. These metrics can be combined to create a composite measure, referred to here as the validation score. The validation score is used to compare results obtained across a range of values for one or more sorting parameters.

Metrics to evaluate sorting quality have been developed previously (Fee, Mitra, & Kleinfeld, 1996; Harris, Hirase, Leinekugel, Henze, & Buzsáki, 2001; Joshua, Elias, Levine, & Bergman, 2007; Pouzat, Mazor, & Laurent, 2002). Most focus on only a single statistical attribute of the data, such as cluster isolation or apply only to a particular sorting algorithm. Given the limitations of using only statistical metrics, many users prefer manual waveform inspection to determine quality (Hill et al., 2011; Schmitzer-Torbert, Jackson, Henze, Harris, & Redish, 2005), which is highly subjective, time intensive, and prone to human error and variability. Our heuristic metrics evaluate many different parameters of the data, inspired by the types of features that human evaluators typically identify, which are in turn motivated by neuroscientific and electrophysiological principles of extracellular recording.

We propose the Heuristic Spike Sorting Tuner (HSST) as a framework to maximize sort quality and avoid the pitfalls of standard methods to enable truly unsupervised spike sorting. Importantly, the parameter tuning provided by the HSST framework is applied to all steps in the spike signal processing chain, to allow iterative tuning of parameters affecting spike detection, feature extraction, and clustering.

Experienced users of spike-sorting software will recognize that many algorithms may work well for one specific dataset but may perform poorly for another. HSST avoids this pitfall, by parallel sorting a dataset (across a range of parameters) and using a validation score to select the best output. A set of heuristic metrics (which explore individual feature spaces of the extracted waveform snippets) test each output, scoring the sorted result based on a set of focused criteria. HSST combines the results of these many metrics to build a strong classifier, and is able to determine the best output of a sorting algorithm without any human input.

## METHODS AND MATERIALS

Spike sorting is frequently approached as a multi-step process and is broken down into three steps: 1) spike detection, 2) feature extraction, and 3) clustering (Gibson et al., 2008). The HSST framework adds a fourth step: parameter validation and selection. For example, spike waveforms are often detected by setting an amplitude threshold and extracting a short segment of the continuous data surrounding the threshold crossing event. Next, a feature extraction process is used to identify attributes of the spike waveforms that are uniquely associated with individual neurons. These features or attributes can include amplitude measures (e.g. peak to peak, positive or negative peaks), but a common approach is to apply principal components analysis (PCA) (Pearson, 1901) or other dimensionality reduction methods to extract a low dimensional representation of the spike waveforms that capture the discriminable features. Finally, the process of sorting the spike waveforms (Table 1) into different “units” is performed by partitioning the spike features into separable groups using clustering algorithms. HSST provides a framework for auto-tuning parameters in spike detection and sorting algorithms.

**Table 1:**
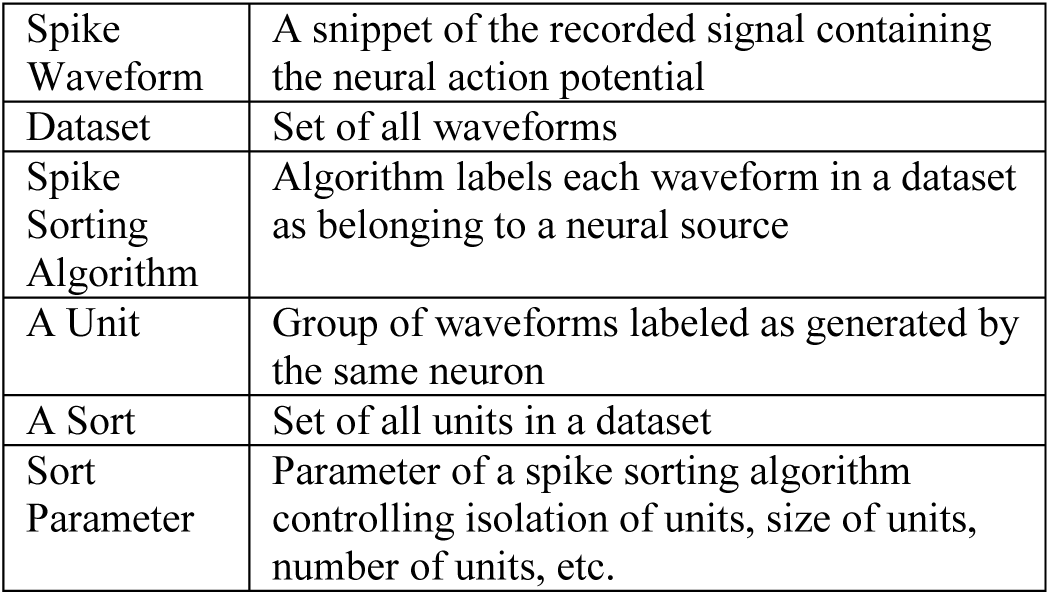
Spike sorting definitions used throughout this paper.

### Overview of the HSST Framework

The primary objective of the HSST is to enable sorting algorithms to correctly identify well-isolated spikes. To best accomplish this, a parallel algorithm (Figure 1) is used to detect the best parameter set after sorting, rather than the traditional way of estimating the best set of parameters, prior to sorting. The framework sorts a dataset N times with the user’s preferred sorting algorithm. Each time, it uses a different set of parameters or initial conditions, yielding *N* results. The framework has a fixed cost function to score each resulting sort, selecting the output which best groups detected spikes into distinguishable clusters.

**Figure 1:**
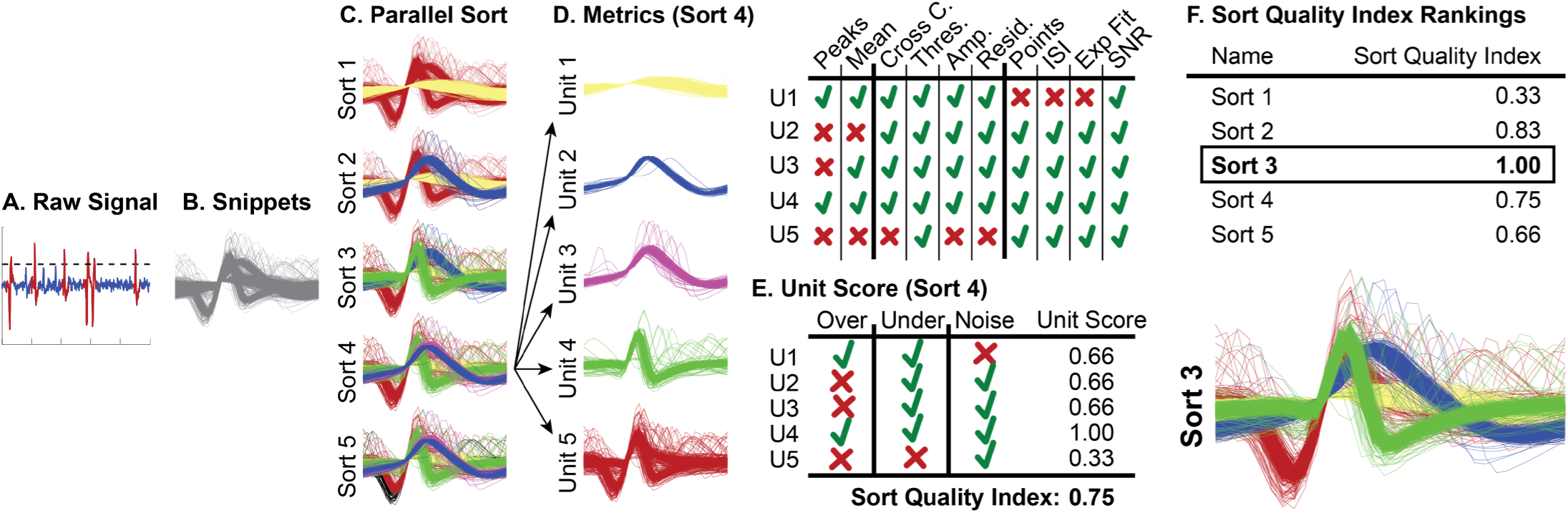
HSST flow diagram for sorting a particular dataset. (A-B) The raw electrical signal is thresholded to extract snippets. (C) Snippets are sorted using a given sorting algorithm using a range of parameters. Each parameter set yields a unique sorting result, Sorts 1-5 in this example. (D) The units identified in each sort are assigned a score based on each metric. Sort 4 is scored as an example. Green check marks mean a unit passed this particular metric, i.e. the absence of a particular error type. Red crosses mean an error type was found. Based of those results, each unit is labeled either over-sorted, under-sorted or noise (E). The Sort Score is the average of each of the unit scores, weighted by the number of waveforms in each unit. Each Sort Score, called the Sort Quality Index (SQI), is ranked. The highest scoring sort is selected as the output of HSST.

### Feature Spaces

The HSST framework uses a set of feature spaces derived from the broadband neural data. The features are broadly separable into two groups: amplitude features and temporal features. The amplitude features include:

1. Minimum and maximum amplitude of the spike waveform
2. Distribution of waveform amplitudes within a unit
3. Distribution of the derivatives of the waveform
4. Similarity of a waveform to the unit’s mean waveform
5. Similarity of a waveform to another unit’s mean waveform
6. Number of inflection points in a unit’s mean waveform

The temporal features include:

1. Distribution of inter-spike-intervals (ISI)
2. Location of a unit’s min/max peaks across units

### Heuristic Metrics

The strength of the HSST framework lies in considering a large set of metrics each contributing a vote, generating a strong, robust classifier. Ten heuristic metrics are used to identify potential sorting errors that we grouped into three Error Categories (Table 2); over-sorted errors, under-sorted errors, and noise errors.

**Table 2:** HSST definitions of evaluated unit quality. Each unit can be labeled as any combination of the three error categories. If any metric (right most column) in each category fails, the unit being sorted is labeled with that error category.

| Error Categories | Definition | Metric Name |
| --- | --- | --- |
| Over-sorted | A unit which contains some waveforms from a source neuron, but fails to contain all of them | Dissimilar Peaks |
|  |  | Mean Waveform |
| Under-sorted | A unit which contains waveforms from multiple source neurons | Cross Correlation |
|  |  | Threshold Slope |
|  |  | Max Amplitude |
|  |  | Residuals |
| Noise | A unit which contains multi-neuron activity, recording artifacts, electrical noise, etc | Stationary Point |
|  |  | ISI Violations |
|  |  | Exponential Fit |
|  |  | Low SNR |
|  |  | Under-Sorted |

These metrics and categories were selected based on the goal of identifying features of waveforms that should be sortable, rather than trying to tease apart low amplitude signals where it is often difficult or impossible to distinguish one neuron from another. For a unit containing waveforms with an SNR that should be able to be sorted, there can be both under-sorting errors as well as over-sorting errors. An over-sorting error occurs when a unit contains waveforms from just a single neuron, but not all of the waveforms from that neuron. An under-sorting error occurs when a unit has waveforms from multiple neurons. Finally, a noise error occurs when a unit contains waveforms with low SNR that represent multi-unit activity or other noise artifacts.

The eleven metrics vote *pass* or *fail* on each unit in a sort (Figure 1D). A *pass* means a particular metric did not find a particular error type in a unit, while a *fail* means an error type is found. See the section on metrics below for details on how each is tabulated.

### Unit Scores

Each unit is assigned a “Unit Score” (Figure 1E) based on the criteria outlined in Table 2. The Unit Score ranges between 0 and 1; the higher the unit score, the more likely that unit contains spike waveforms from a single neural source. Each unit begins with a perfect score of 1 (a well sorted unit) and loses 1/3rd of a point for each type of sorting error present in the unit (over-sorted, under-sorted and/or noise).

### Sort Quality Index (SQI)

The sort quality index (SQI) is the ranking used to determine which sort has best grouped waveforms according to their source neuron. By maximizing the SQI, we can generate the best overall performance of a particular algorithm. The parameter set with the highest SQIs are the most likely to have correctly labeled each waveform with its true neural source.

The SQI is calculated as the average of each of the unit scores within a sort (Figure 1E), weighted by the number of waveforms in each unit. The value can range from 1 (all units contain the activity of a single source neuron) to 0 (units have waveforms from a mixture of multiple source neurons and/or noise).

### Illustration of Framework Logic and Function

Using a computationally generated dataset (see Datasets section for full details), we demonstrate the full breadth of calculations used to determine the optimal sorting output (Figure 1). Our initial dataset contains three neurons in close vicinity to the electrode (50 µm), with multiple other neurons further away (>200 µm). The dataset is sorted with K-Means five times, each with a different parameter, K = 2, 3, 4, 5, 6. The unit scores for each sort are ranked (Figure 1D) and averaged (Figure 1E) to give a sort quality index (SQI). Each unit score is normalized by the number of waveforms in that unit. If a single unit is identified as noise (the yellow unit in Figure 1D), it is not included in the averaging to obtain the final score. It is desirable to group any noise waveforms together and isolate them from the rest of the dataset, therefore the largest noise unit is not counted in the final average, but any additional noise units are. We can see that Sort 3 yields a result most resembling the ground truth. HSST scores this sort the highest SQI (Figure 1F).

### Metrics for Detecting Over-sorting Errors

Over-sorted units are units where the waveforms from a single neuron are split into two or more units. Intuitively, this means that waveforms from a single source neuron are incorrectly labeled as two or more distinct neural sources. Over-sorting will cause underestimation of neuronal firing rates and could significantly modify calculated firing patterns.

The output of each of these metrics is binary: pass or fail; if any over-sorting metric detects an error, then the unit fails and is labeled over-sorted. If the unit passes all over-sorting metrics individually, meaning each metric verifies that no error type exists in the unit, then those waveforms are appropriately sorted and do not overlap with other unit’s waveforms. These metrics are only used if multiple units are identified in a given sort. If only a single unit is identified, these metrics all pass.

#### Dissimilar Peaks

If the waveforms of a single neuron are separated into two different units, we would expect the distributions of the positive and negative waveform extrema for those units to be similar. For this metric, all units are pairwise compared. The mean and standard deviation for both the positive and negative waveform values is first calculated for each unit. The D’ statistic (Vallbo & Johansson, 1966) is then used to compute the D’ distance between the positive and negative extrema for all pairs of units (Figure 2B,D). The D’ statistic ranges from 0 to positive infinity, and a value less than one indicates statistically significant overlap. Three conditions are required for a pair of units to be identified as over-sorted using this approach: 1) the positive extrema distributions must have a D’ distance less than one, 2) the negative extrema distributions must have a D’ distances less than one, and 3) the overlapping extrema fall within 0.125 milliseconds (2-3 sample points) of each other. The third condition places a reasonable temporal constraint on the timing of the positive and negative extrema. Any pair of units that meet all three criteria are considered to be over-sorted and fail this metric. If any of the conditions are unmet, the metric passes, and thus are well-sorted.

**Figure 2:**
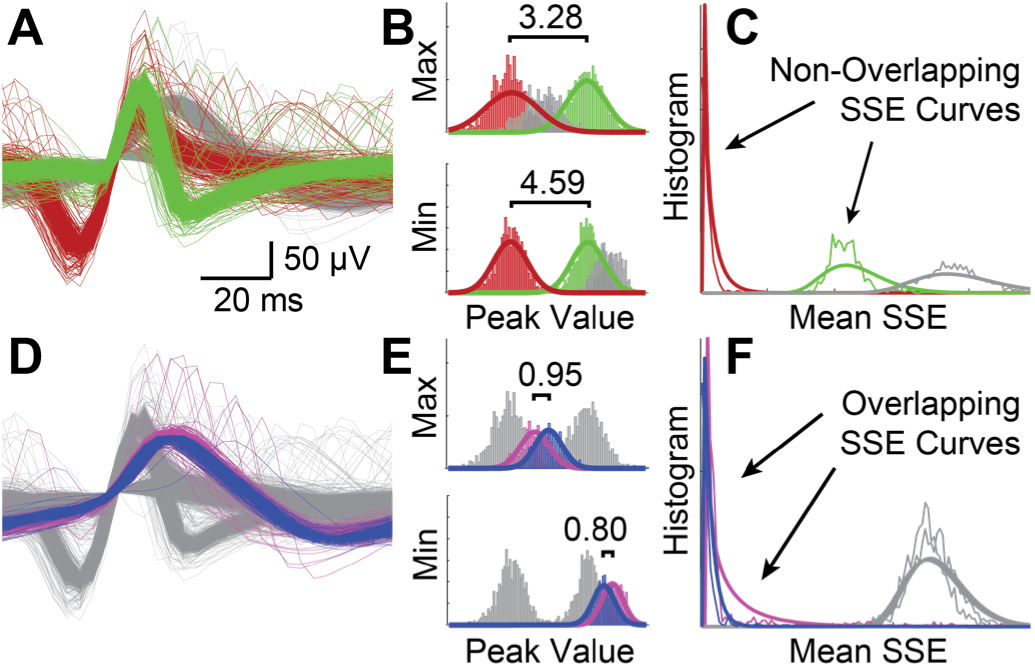
Over-Sorted Metrics. Examples of two well-sorted units (A), shown in green and red, in the waveform space. The D’ distance (or sensitivity index) is used to evaluate the overlap between the positive/negative extrema distributions between all units pairwise in a given sort (B). We can also compute a distribution of the sum of squared errors between the mean waveform and each waveform in the unit (C). For two well-sorted units, these distributions do not overlap. We also show examples of two over-sorted units (D) in the waveform space. Analogous to our analysis in (B) and (C), we see these units on the bottom overlap significantly in both feature spaces (D,E).

#### Mean Waveform Sum of Squared Errors

In order to compare waveform shapes as a whole (instead of only the extrema), we calculate two distributions: 1) the sum of squared errors between the mean waveform of a specific unit and all the individual waveforms found in that unit, and 2) the sum of squared errors between the mean waveform of that unit and all the waveforms of a second unit. These two distributions are combined and multi-modality is assessed (Figure 2C,F). If the combined distribution is unimodal, the unit is identified as over-sorted and fails this metric. This process is iteratively applied to each unit, such that every unit is compared to every other unit in the sort group. If no overlap with any other unit is found, then the metric passes.

### Metrics for Detecting Under-sorting Errors

An extracellular recording of an action potential from each source neuron will produce a stereotypical waveform shape. The distribution of waveform shapes recorded from a single source neuron can be approximated by a mean waveform with some mean-zero Gaussian distribution at each sample point. However, an under-sorted unit–a unit with waveforms from more than one source neuron–will violate this assumption of mean-zero Gaussian distribution. With these assumptions, we can test for unimodality along different dimensions in the waveform feature space to examine whether multiple neural sources are present within a single unit. Similarly to the previous category of metrics, the output of each of these metrics is pass or fail. If any one of these metrics detects an error type, the unit fails and is deemed under-sorted (has multiple neural sources). If all metrics pass, by verifying no error present, then the unit is well sorted (only one neural source).

These metrics rely on determining the number of modes (or statistically separable groups) in some feature space of the data. Many other techniques (Hartigan & Hartigan, 1985; Rozal & Hartigan, 1994; Sawitzki, 1996) have been developed to detect multi-modality, but due to the variability of neural data, we developed our own standard way of detecting multi-modality (Appendix, Figure A1).

Four different waveform feature spaces are analyzed for multi-modality using the methods described above. Each feature space is used to compute a distribution; checking each for multi-modality constitutes a different metric. Below we describe calculations to compute each distribution.

#### Residuals

We compute a measure of residuals by calculating the mean waveform of the unit and the variance about that average waveform (Figure A3). At each time point, the distribution across all waveforms in the unit is checked for multi-modality (Figure 3B). In addition, if the measured variance of that distribution lays outside the estimated standard deviation, the unit is considered multi-modal.

**Figure 3:**
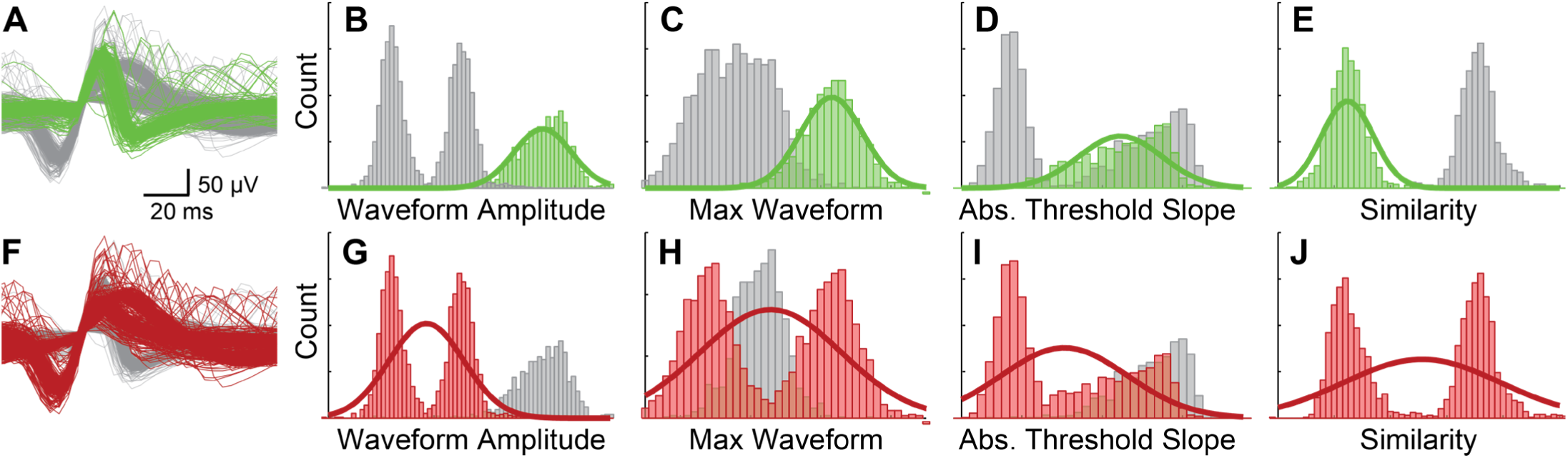
Under-Sorted Metrics. (A) A well-sorted unit is analyzed for multi-modality in four different waveform features spaces (B-E). (B) The waveform amplitude metric examines a slice of each waveform amplitude at each time point. (C) The maximum amplitude metric looks at the max amplitude of each waveform. (D) The threshold slope metric examines the slope of the waveform as it crosses the threshold. (E) Similarity metric between each waveform and the mean waveform of the unit. (F) The under-sorted unit on the bottom has examples of each waveform feature space for each metric (G-J). Note that for each metric, the green distributions a well-sorted unit) are unimodal. However, the under-sorted unit, which contains waveforms from multiple different neurons has a bimodal distribution for each metric.

#### Peak-Waveform Amplitude

We calculate the maximum amplitude of each waveform in the unit and check that distribution for multi-modality (Figure 3C).

#### Threshold Slope

We calculate the slope of each waveform as it crosses the threshold value (Figure 3D). This distribution is checked for multi-modality.

#### Waveform Similarity

We compute a waveform similarity measure by calculating the cross correlation between the mean waveform of the unit and each individual waveform in that same unit. The maximum value of each cross-correlation is the similarity between that individual waveform and the mean waveform (Jackson & Fetz, 2013). We check for multi-modality in a histogram of similarity from each waveform (Figure 3E).

### Metrics for Detecting Noise Errors

Low SNR neural recordings are common, especially on multichannel microelectrode arrays. These waveforms are typically the most challenging to sort. In many research applications, it is beneficial to collect all the noise waveforms into a single unit labeled as noise or multi-unit activity. Therefore, we might expect that if the spike detection threshold was low, such a noise unit would exist. The purpose of the metrics described below is to detect and isolate those noise waveforms from the rest of the units. The output of each of these metrics is a pass or fail; if any of the noise metrics detects an error type, the unit poorly sorted and contains noise waveforms. However, if all metrics verify no error present, the unit passes and does not contain noise waveforms. Below we detail each metric’s algorithm individually.

#### Inter-Spike Intervals (ISI)

A count of inter-spike interval (ISI) refractory violations is a reliable metric for a poorly isolated unit. Biologically, all neurons have absolute refractory periods of approximately one millisecond, meaning that a neuron cannot fire at a rate greater than 1000 spikes per second. In practice, neuronal firing rates are much lower, with the fastest neurons peaking at several hundred spikes per second (Gittis, Moghadam, & Lac, 2010). For our study with generated data, we assumed our generated neural activity fired a random rate less than 1 kHz. If more than 5% of the ISI measured were less than 1 millisecond, the unit is identified as noise (Figure 4C,F).

**Figure 4:**
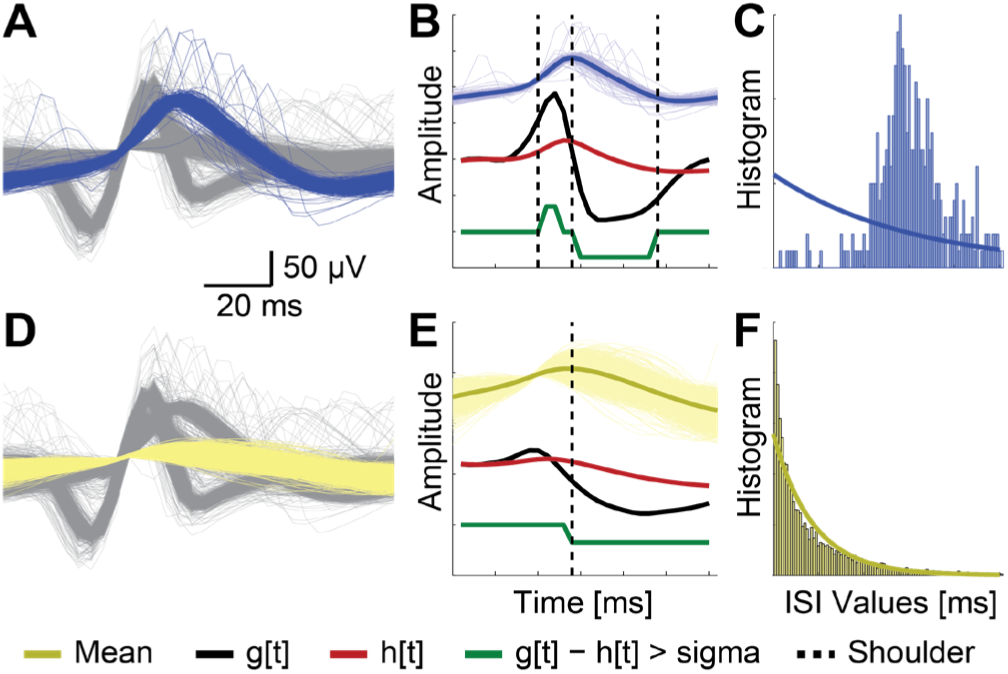
Metrics for Determining Noise. Example waveforms of a well-sorted unit (A) and a noise unit (D). Shoulder points (B) can be identified by the threshold crossings of the two derivative curves (black and red) of the mean waveform. The red derivative, g[t], is slope between sample point t and sample point t+1. The black derivative, h[t], is the slope between sample point 1, and sample point t. The green line shows inflection which occur when the difference between the red and black lines is greater than one sigma. Shoulder points are identified by dotted vertical lines. A histogram of Inter-Spike-Intervals (ISI) and an exponential fit (C). Since the fit is poor, this unit is not considered a noise unit. The exponential fit to the ISI distribution in F shows a good fit, indicating that this unit is likely a noise unit.

#### Single Stationary Point

Extracellular recordings of neural spiking activity tend to exhibit bi- or tri-phasic spike waveforms. If we identify a unit with a single “bend” or “shoulder” in the mean waveform, it is likely to be a noise unit. We identify shoulders in the mean waveform by examining the crossing points of two curves (Figure 4B). The first curve is the derivative of the mean waveform. The second curve is the slope from the first sample point of the waveform to the nth sample point. Not all crossing points are significant however. The standard deviation of the absolute difference between the two curves is calculated. Only crossing points with at least two consecutive points between them which have a difference greater than one standard deviation are counted in the total. If one or zero “bends” are detected, then the unit is identified as noise (Figure 4B,E).

#### Inter-Spike-Interval Exponential Fit

If we model the background multi-unit activity or electrical noise as a stochastic mathematical process, for example a normal random process, we can detect noise units by identifying the characteristics of noise generated from our model. When we set the spike threshold to an arbitrary value, and “detect” spikes from data generated from this model, the histogram of times between the points above that threshold, or inter-spike intervals (ISI), can be approximated by an exponential curve. Therefore, if a unit’s ISI histogram fits an exponential curve with a normalized mean squared error (NMSE) rate of 15% or less, this unit is categorized as noise (Figure 4C,F).

#### Signal-to-Noise Ratio (SNR)

Keeping with HSST’s primary objective to enable sorting algorithms to correctly identify well-isolated spikes, we therefore introduce a final, optional metric to eliminate low SNR units. Units whose mean waveform are below an SNR of 1, fail this metric and are labeled as noise.

We use an iterative process to calculate SNR from the raw waveform (Figure A2). Signal are considered data points greater than four standard deviations from the mean of the raw waveform. These data points are removed and a new standard deviation is calculated from the remaining data. This process is repeated until the estimate of standard deviation converges (Figure A2). One standard deviation of the leftover data of the final round becomes our estimate of noise. If the raw waveform is not provided, the user will need to provide an estimate of the noise. (Again, this metric is optional to help users looking to get only high SNR spikes. Users can also manually determine this SNR threshold for their particular need if so desired).

### Datasets

In order to accurately compare the performance of spike sorting algorithms, it is necessary to know the ground truth regarding the identity of each recorded spike. In experimental electrophysiological recordings, the ground truth is unknown, so we used a computational simulation to generate test datasets. The NeuroCube (Camuñas-Mesa et al 2013) model was used to simulate a recording electrode in a volume of conductive tissue containing electrically active neurons. This model includes an electrode as a single point in a 1 mm cube of conductive tissue, containing several neurons 50-200 µm away from the electrode as well as hundreds of background neurons (> 200 µm away). To compare performance results between different sorting algorithms, we used the same dataset used for testing the WaveClus algorithm (Quiroga et al., 2004). This dataset (Figure 5) contained 4 groups, each with 3 neurons in close proximity to the simulated recording electrode, and hundreds of neurons further away to simulate background neural noise. Two groups contained three neurons with very similar waveform shapes (Figure 5A,B) while two groups contained neurons with dissimilar waveform shapes (Figure 5C,D).

**Figure 5:**
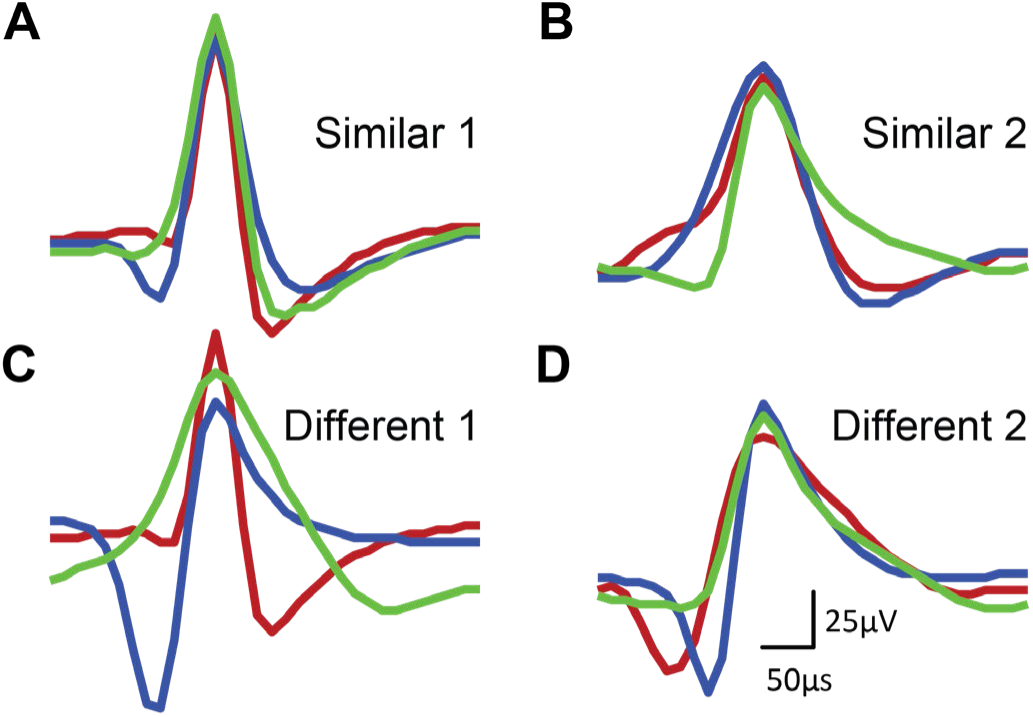
The mean waveform from each source neuron (shown in red, blue and green) from the 4 groups in the WaveClus dataset (Similar Shape 1 & 2, Different Shape 1 & 2) is shown.

In order to understand the effects of SNR on sorting accuracy, we tested on each group at four different distances from the electrode (50 µm, 100 µm, 150 µm, and 200 µm), generated by the model defined in Equation A1. This resulted in testing each algorithm 16 times, on each of the four groups at four distances.

To test sorting accuracy on datasets with different numbers of close proximity neurons, we ran additional tests using the four groups at a fixed distance of 150 µm. We varied the number of close proximity neurons by removing neural waveforms from each dataset. One set had the original 3 source neurons (plus noise), another set 2 neurons (plus noise), and the final set, 1 source neuron (plus noise). This resulted in testing each algorithm 12 times, each of the 4 groups with the 3 different sets of neurons.

The HSST framework was also tested on real neural data consisting of 80 datasets recorded from microelectrodes implanted in the dorsal root ganglion (DRG) of a cat’s L6 and L7 spinal roots (Debnath et al., 2014). For these real world datasets, no ground truth existed; thus, expert human sorters identified all units with an SNR greater than 5, calculated as the ratio of peak signal voltage to estimated peak noise voltage. High SNR units were used exclusively so that just those waveforms that could be conclusively identified as originating from the same neuron were compared. Humans are typically good at identifying units with high SNR, giving us reasonable confidence about the accuracy of these unit identifications.

### Sorting Algorithms

HSST is fundamentally an unsupervised tool to enable existing sorting algorithms to choose their free parameters. We used implementations of six widely used sorting algorithms to compare their performance and evaluate HSST at selecting their correct parameters. GMM is a Gaussian mixture model with a variable number of clusters (Neal & Hinton, 1999). KM is a K-Means algorithm (MacQueen, 1967). WaveClus is the algorithm proposed by Quiroga et al., 2004. DukeSort is the method proposed by Chen et al., 2011 and M-Sorter is the method proposed by Yuan et al., 2012. UltraMegaSort2000 is the method proposed by Hill et al 2011. Code was gathered from public resources provided by the authors while GMM and KM are MATLAB implementations.

## RESULTS

### HSST Parameter Selection

The HSST framework operates by sorting a dataset with a specific algorithm across range of input parameters and selects the optimal sort. For generated data sets used in the following analysis, classification error for these optimal sorts was measured by the number of waveforms assigned incorrectly to the wrong neuron. The classification error varied with distance from the recording electrode (Figure 6) and therefore, the corresponding SNRs. The Gaussian Mixture Model (GMM) algorithm sorted the data with a range of 6 different parameter sets, each yielding a different classification error. The optimal parameter selection made by HSST was compared to the classification error to judge the accuracy of the HSST algorithm.

**Figure 6:**
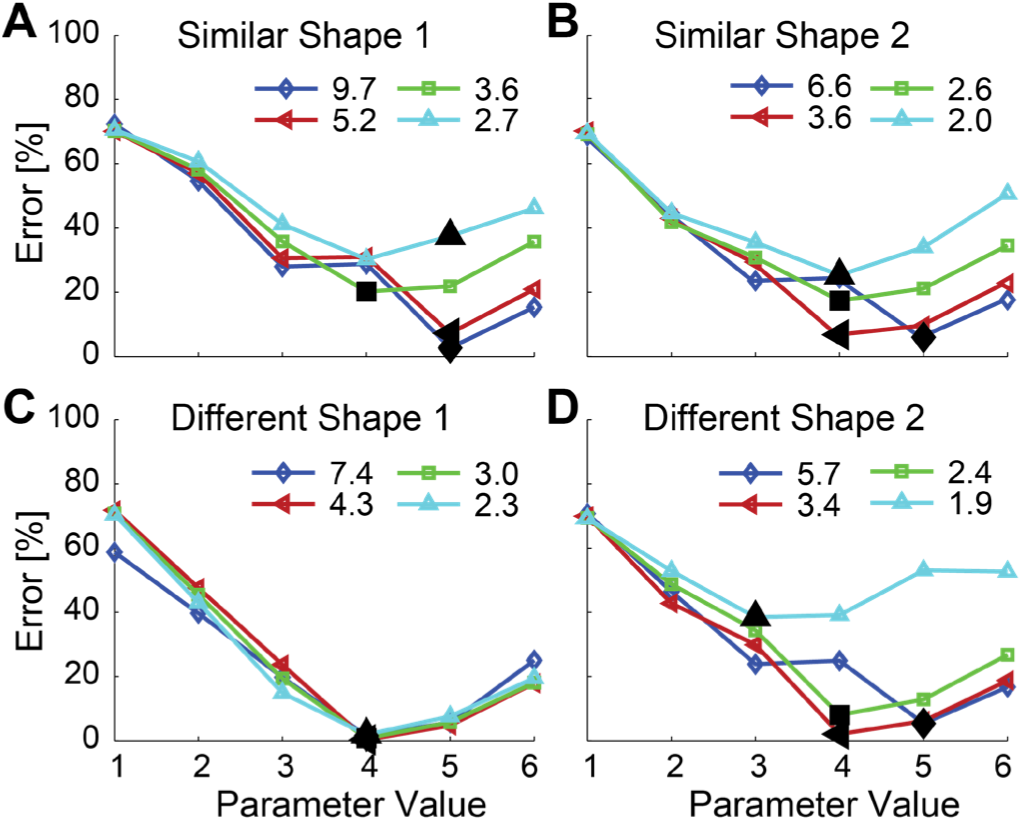
HSST Parameter Selection Error. On each plot, classification error is shown on the Y axis, and the parameter setting is shown on the X axis. The colored lines indicate measured SNRs, corresponding to the distance from the recording electrode (the lowest SNR corresponding to distance 200 um, while the highest SNR corresponds to distance of 50 um). The Gaussian Mixture Model (GMM) algorithm was used to generate the sorts for each data set. The main parameter of the GMM algorithm sets the number of clusters. The Parameter Value in the above plots correspond to the number of clusters. The black markers indicate the parameter setting chosen by the HSST algorithm. We can see that in nearly every case, HSST selects the parameter set with the least classification error.

### Parameter Selection and Poor Sorting Errors

When determining the optimal sorting parameters for a given algorithm, we defined two types of errors, selection error and sorting error, to test the accuracy of HSST.

Parameter selection errors occur when HSST selected a parameter which didn’t minimize the classification error (i.e. selects 4 clusters when only three are present). Poor Sorting errors occurred when the sorting algorithm itself poorly grouped the waveforms, and all outputs are poor (Figure A4).

As an example of selection error, in Figure 6A, the light blue line shows a dataset sorted by GMM using six different parameter sets. Parameter set 4 yielded the lowest classification error; however, HSST chose parameter set 5, because two units were considered too similar when sorting with parameter set 4. Due to low SNR, the distributions of waveform shape of the two neurons overlapped, making it difficult to distinguish between the two units. This selection is an error, because parameter set four yielded the lowest classification error.

Poor sorting errors occurred when the sorting algorithm fell into a local minimum, as illustrated in Figure A4. Here, four clusters were present, yet the algorithm divided the noise cluster into three pieces and combined all three source units. We generated the data therefore the correct number of clusters is known to be four; however, the classification error was high for this parameter. This is an outcome desirable to avoid. Unless the user can account for them and predict when they will occur, these errors can render a sorting algorithm useless. With this in mind, HSST is capable of producing optimal sorts, despite with these errors.

As an example, in Figure 6B on the dark blue line, parameter set 5 yielded the lowest classification error. Parameter set 5 yielded the “wrong” number of clusters, since we generated these data to include three neurons and background noise (totaling four clusters). However, due to a failure of the GMM algorithm, parameter set 4 yielded a non-optimal result (Figure A4). Even still HSST selected the parameter set which minimizes classification error (set 5).

This example illustrates the strength of HSST. Even if the theoretical optimal parameter set is known (in this case four clusters are known to be present) a sorting algorithm may not reach an optimal sort output. Unless a sorting algorithm is guaranteed to reach the global minimum (very difficult to prove and not true for the vast majority of current sorting algorithms), it still could fall into a local minimum, resulting in a poor classification result.

As shown in Figure 7, parameter selection error of the HSST framework had a minimal impact. Frequency and classification error is plotted for each of the 16 generated datasets sorted for all sorting algorithms. Only the cases with differences greater than 0 (meaning HSST did not select the optimal parameter) are shown. When we used HSST in combination with the GMM sorting algorithm, the non-optimal parameter set was only selected one time out of 16 datasets, resulting in an increase of just 8% classification error above the absolute minimum error achievable with the optimal parameter set for that instance. This data point corresponds to the light blue curve in Figure 6. For 90% of all cases, HSST selected either the optimal parameter, or a parameter within 10% classification error of the optimal parameter.

**Figure 7:**
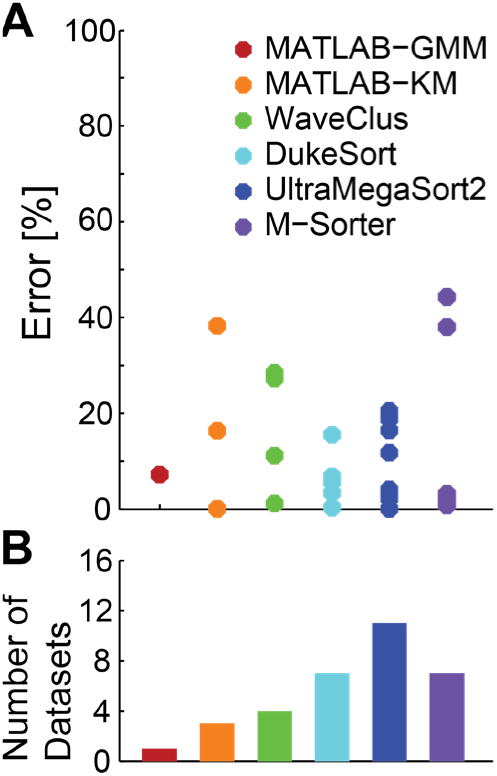
Frequency and Size of Alpha Error Occurrences. (A) Percentage of parameter selection error caused by HSST selecting non-optimal parameter by sorting algorithm. (B) Shows the number of points in *A* in each row (one row for each sorting algorithm).

### Classification error across SNR

In Figure 8, the WaveClus (Quiroga et al., 2004) datasets were sorted using the six sorting algorithms discussed in the methods (each using HSST to select their optimal parameter sets) and classification error rates were compared.

**Figure 8:**
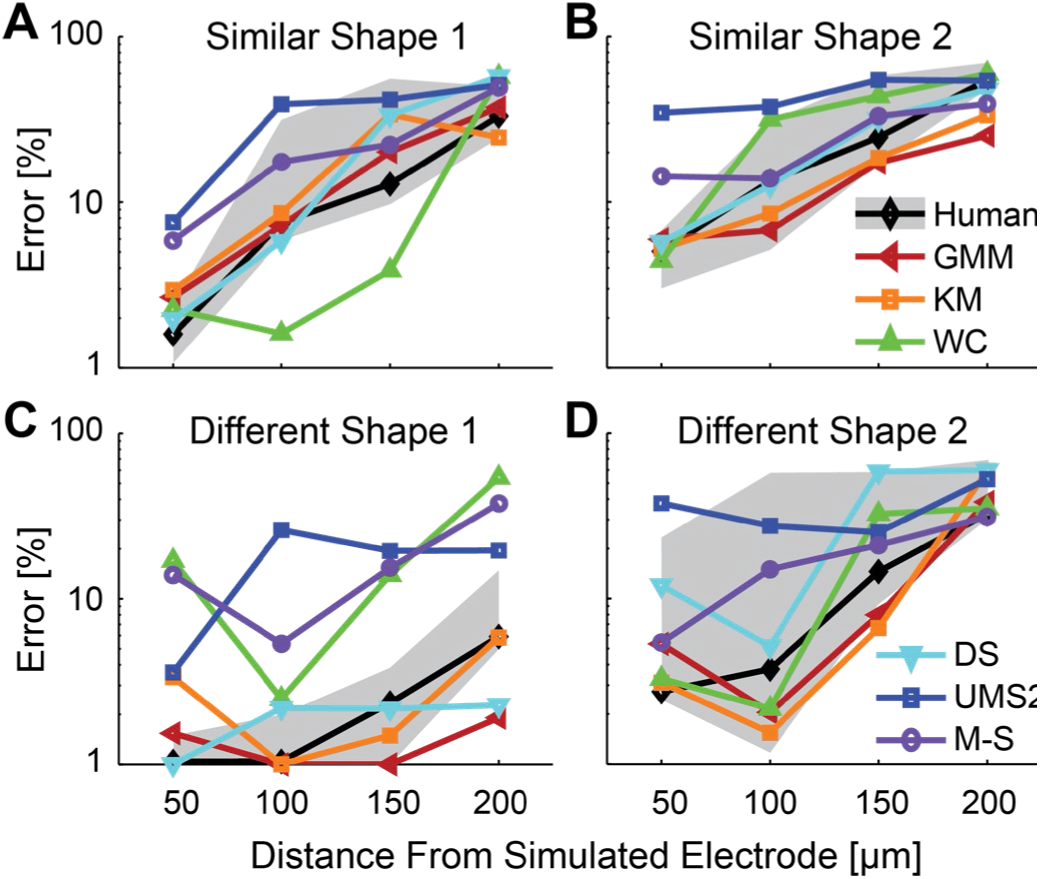
Classification Error. For each of the four datasets, 4 different values of SNR where sorted by each method. HSST sorted the data with each of the different methods across a wide range of parameters. Each of these algorithms (excluding WaveClus) required extensive parameter selection, with high sensitivity in accuracy to selection. HSST chose the best parameter from the given set. Error Bars denote expert human sorter variability. On each plot, classification error is shown on the Y axis, and SNR value of that dataset is shown on the X axis. Each dataset was injected with noise to produce four different SNR values on the X axis.

Several general trends emerged our analysis. As expected, as the distance from the electrode increases (i.e. SNR decreases), performance degrades. Several human experts manually sorted the same datasets, and their average performance is shown in black with grey error areas. The error areas show the max/min values of the human sorters. Surprisingly, simple algorithms such as K-Means and GMM performed just as well as humans and often outperformed the other more complex algorithms when combined with HSST.

We analyzed the sensitivity of each algorithm to changes in SNR (proportional to distance between neuron and recording electrode), and the sensitivity of the algorithms to changes in the range of parameters (Figure 9A). As shown in the grey boxes, we showed the classification error produced by the range of parameters given to each of the algorithms (Figure 9A) This range was calculated by selecting each of the input parameter sets and evaluating the resulting classification error. The colored boxes show the range of error of only the optimal parameters selected by HSST. For example, the range of input parameters given to GMM yielded a large range of error regardless of SNR. However, the range of parameter selected by HSST produced low classification error. On the other hand, for UltraMegaSort, the range of error yielded by all parameters is much tighter. This algorithm is not as sensitive to parameter selection, and therefore doesn’t benefit as much from HSST selecting its parameters.

**Figure 9:**
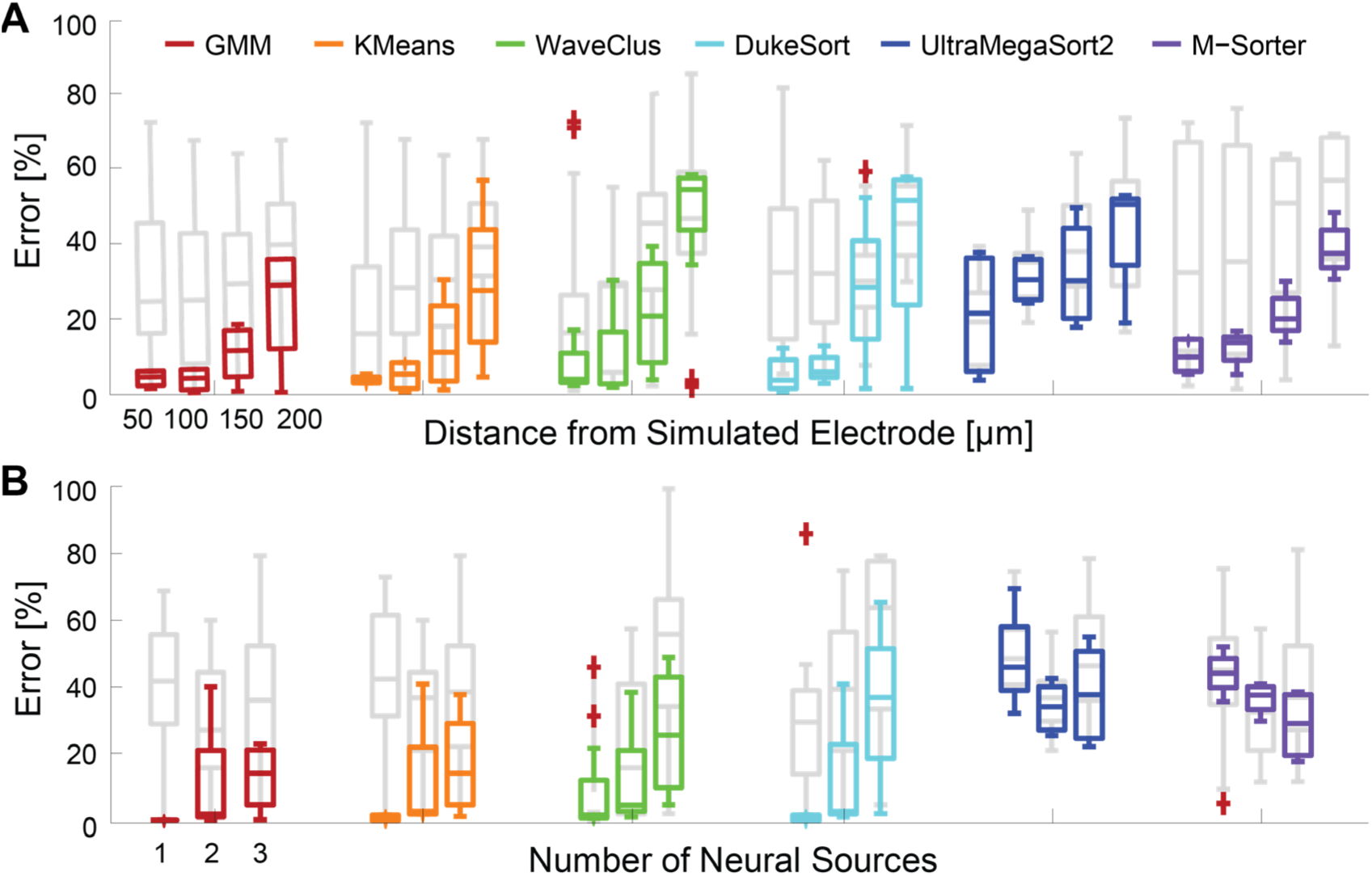
Parameter Sensitivity to Changes in SNR (A). Boxplot showing range of error resulting from parameter selection. Colored boxes show range of error from HSST selection. Grey boxes show range of error from all parameters included in the sweep. Some algorithms have high sensitivity to changes in SNR, other show little change in variability of error. Parameter Sensitivity to Changes in Number of Units (B). Modifying the number of clusters showed similar results to modifying SNR. As the number of clusters decreased, we saw improved performance on nearly every sorting algorithm. Simulated distance from the electrode was 150 µm.

### Variability in Number of Units

We also analyized the performance sensitivity to changes of the number of proximal neurons (Figure 9B), while SNR was held constant. As the number of neural sources decreased, identifying the number of clusters became much easier as evidenced by a decrease in classification error. We plotted the parameter range error (shown in grey) and the range of error of parameters selected by HSST (shown in color by sorting algorithm).

### Comparison to WaveClus (fully unsupervised version)

To validate performance of the HSST algorithm against another unsupervised algorithm, we compared its accuracy against performance of an expert human sorter and an additional package in WaveClus codebase (Quiroga et al., 2004). This additional package has an automated method for estimating the two free parameters (temperature and minimum cluster size) of the Waveclus algorithm, making it a fully unsupervised. HSST’s performance in classifying waveforms was at least as good as a human and outperformed WaveClus (Figure 10).

**Figure 10:**
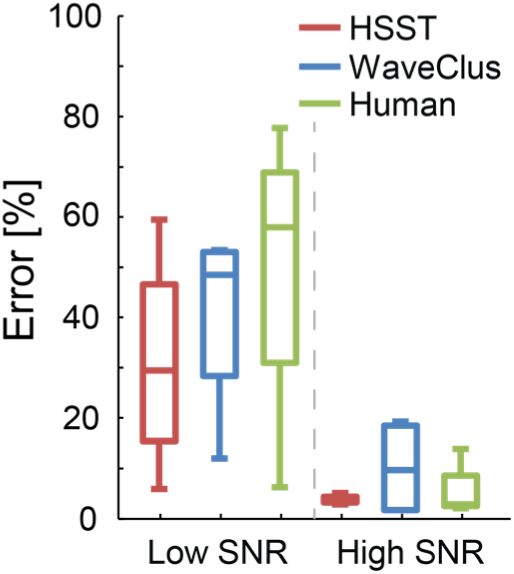
Average classification error between various sort methods on the generated data. Results are an average of the four datasets in Figure 11 (highest and lowest SNR values only) compared to average error of four expert human sorters. Because we are averaging across different datasets, the SNR values are approximately 2 (for the low SNR value) and 5 (the high SNR value). We see that the HSST (with a simple K-means sorting algorithm) can do at least as well as a human and does better than the other algorithms. Graph shows a boxplot, in which the middle bar shows the median value of the data, edges of the box show quartiles (25% and 75%) and whiskers show most extreme data points excluding outliners.

### Real Neural Data

To demonstrate performance on real neural data, we sorted data recorded from the cat dorsal root ganglia (DRG), when parameters in the algorithms were set by HSST. Since the ground truth is unknown in this dataset, expert human sorters labeled high SNR waveforms into units. We tested how well HSST could identify these units and label the rest of the waveforms as noise.

Three statistical metrics were used to quantify performance (Figure 11) of these sorting algorithms: accuracy, sensitivity and specificity (Metz, 1978). Accuracy measured the percentage of correctly classified waveforms, either belonging to a high SNR unit or noise. To further understand how the waveforms were grouped by HSST, we plotted sensitivity (the likelihood of correctly labeling a waveform as a high SNR signal) and specificity (likelihood of incorrectly classifying a noise waveform as neural).

**Figure 11:**
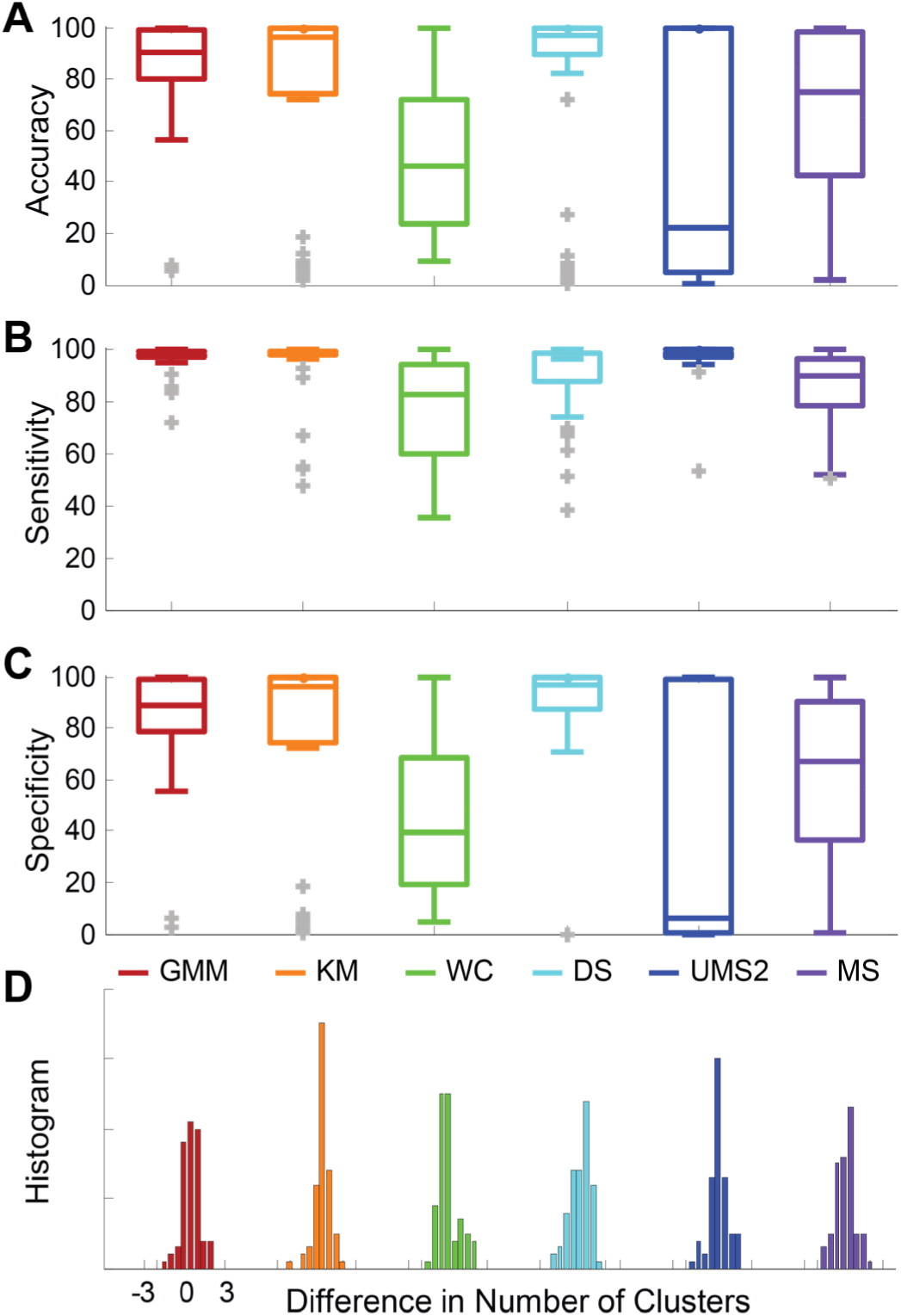
DRG Sorting Results – (A) Accuracy, (B) Sensitivity, (C) Specificity. 80 datasets (classified by an expert human sorter, each with one high SNR unit) were sorted using six sort methods (all of whom had HSST selecting their free parameters). (D) The difference between the total number of clusters identified by the expert human sorters and the number of clusters identified by the sorting algorithm. Each of these histograms has zero mean with some standard deviation demonstrating no bias towards over or under selecting the number of clusters.

A high accuracy score resulted from correctly labeling many waveforms; however, to analyze the individual failure modes of each algorithm (ie. did error result from missing neural activity or from labeling noise as signal), we needed sensitivity and specificity. Sensitivity measured correct identification of neural signal and specificity measured the rejection of noise. A high sensitivity score meant the classifier correctly labeled high SNR units, while a low sensitivity score meant the classifier incorrectly labeled high SNR waveforms as noise. High specificity scores meant noise waveforms were not included in high SNR units.

Eighty datasets (each with one high SNR unit classified by an expert human sorter) were sorted using the six sorting algorithms compared previously (all of whom had HSST selecting their free parameters) (Figure 11). These results showed K-Means, GMM and DukeSort did the best job of reliably selecting these units, while WaveClus and M-Sorter usually included waveforms which didn’t belong in the cluster and UltraMegaSort2000 usually failed to find the high SNR unit. The bottom plot showed the difference between the total number of clusters identified by the expert human sorters and the number of clusters identified by the sorting algorithm. Each of these has zero mean, indicating HSST isn’t biased towards over or under selecting correct number of clusters.

### Speed and Calculation of HSST Sort Score

To quantify the additional time (not including time for spike detection, feature extraction, or clustering) needed to score each sort result, we used HSST to score sort results for various numbers of waveforms and various numbers of units. Figure A5 showed total run time dominated by the sorting algorithm, while running HSST has very little overhead. Most algorithms have comparable runtimes, except for DukeSort which can take longer. These results show that computationally intensive algorithms do not necessarily translate to better performance.

## DISCUSSION

The most vital aspect of this algorithm is its ability to determine the optimal parameters, given a finite set of parameters, for a specific sorting algorithm for the purpose of generating a reliable and physiologically plausible result. Our main design goal for the HSST was to consistently sort waveforms with high SNR, as these are the most likely signals to be used in further analyses. By using HSST for parameter selection, many sorting algorithms can avoid falling into local minima through over-optimizing on a narrow subset of training data. By sorting with a large range of parameters through HSST, they can achieve good results without requiring significant human oversight when sorting widely varying data (both SNR and number of clusters). By leveraging information known *a priori* about the biological constraints of neural firing (e.g. refractory period, multiphasic action potential waveform shape), we hope to improve the performance of sorting algorithms.

Every sorting algorithm has a set of parameters (whether they are exposed to the end user or not) which determine the results of the algorithm. From the obvious K-Means example where *K* sets the number of clusters, to M-Sorter, which requires the user to specify the minimum distance from the generated template, those parameters are often optimally tuned for each individual dataset. While potentially useful to the user when sorting only a few datasets, these parameters cannot be fixed across a wide variety of data. HSST is designed to account for biological constraints in selecting the optimal parameter values to achieve the best possible sort for a given algorithm.

HSST is not immune to some limitations of parameter fixing, for instance the SNR metric discards low SNR units which are potentially undesirable. However, when discretizing data from a continuous data set, some parameter-fixing is inevitable, and in the case of the SNR metric, if the user does not want to discard low SNR units, they can simply set the SNR threshold rejection threshold to 0. The multi-modality function has several fixed parameters, including the valley depth to determine multi-modality. These values where determined empirically, however, are easily translatable to an accuracy score or some other measure of overlap between two one-dimensional Gaussian distributions. The user can set these parameters accordingly if the results are poor. Each of the parameters fixed in the HSST algorithm were held constant for all analysis done in this paper (Table A1).

The final score is an average of the three category scores, over-sorting, under-sorting, and noise. Some issue could be raised about the decision of how to incorporate each of the different metrics into those three scores. Certainly each of the three error categories could be strengthened by the addition of other metrics; however, there is little data/evidence to suggest a right or wrong way to classify each of these conditions other than evaluating their end result of parameter selection. One could also imagine determining a weighting vector to attach to each of the metrics, depending on the strength of each metric.

Another topic for future study is the potential benefit of an iterative search of the parameter space, explored by HSST. One could argue the parameter space explored by K-Means or GMM is well understood and intuitive, while the more complex algorithms might have unintuitive multi-dimensional parameter spaces (yielding non-monotonic classification error results). A more complex parameter space might cause poor performance, allowing HSST to miss the optimal parameter.

The HSST framework was designed for extensibility, allowing for additional sorting methods, additional metrics exploring new feature spaces, new detection/feature extraction processes or even a new process for searching the parameter space. The heuristic metrics provide the sort quality validation scores which judge the sorted results, allowing HSST to select the optimal parameter set.

The HSST framework is publicly available for use with Matlab 2018a (www.github.com/davidbjanes/HSST). The code for all algorithms tested in this study are also available. Templates for adding additional metrics, sorting algorithms, and detection/feature extraction methods are detailed in a README file.

## APPENDIX Multi-Modality Function

Many of the metrics used to detect under-sorting rely on determining the number of modes (or statistically separable groups) in some feature space of the data. Many papers and other techniques (Hartigan & Hartigan, 1985; Rozal & Hartigan, 1994; Sawitzki, 1996) have been developed to accomplish this task, but due to the variability of the data, a standard way of detecting multi-modal data was developed here (Figure A1), combining several previously developed methods.

Procedure

1. A one dimensional histogram (consisting of N bins) is generated from the data to be analyzed. N is the square-root of the number of data points (F & JW, 1977, p. 49).
2. The histogram is normalized (bin height corresponds to the percentage of total histogram area) and smoothed (each bin value is chosen as the median of itself, the bin immediately preceding and following it (Cocatre-Zilgien & Delcomyn, 1992)).
3. Consecutive bins which are less than 1% of the total area, beginning and ending the range of the histogram, are labeled “insignificant” and removed from analysis.
4. All peak/valley pairs of the smoothed histogram are identified. Their differences in height are normalized between 0 (corresponding to the 1% significant bin area threshold) and 1 (the positive peak amplitude of the histogram).
5. Any peak/valley pairs with a height difference of less than 0.25 are removed.

**Figure A1:**
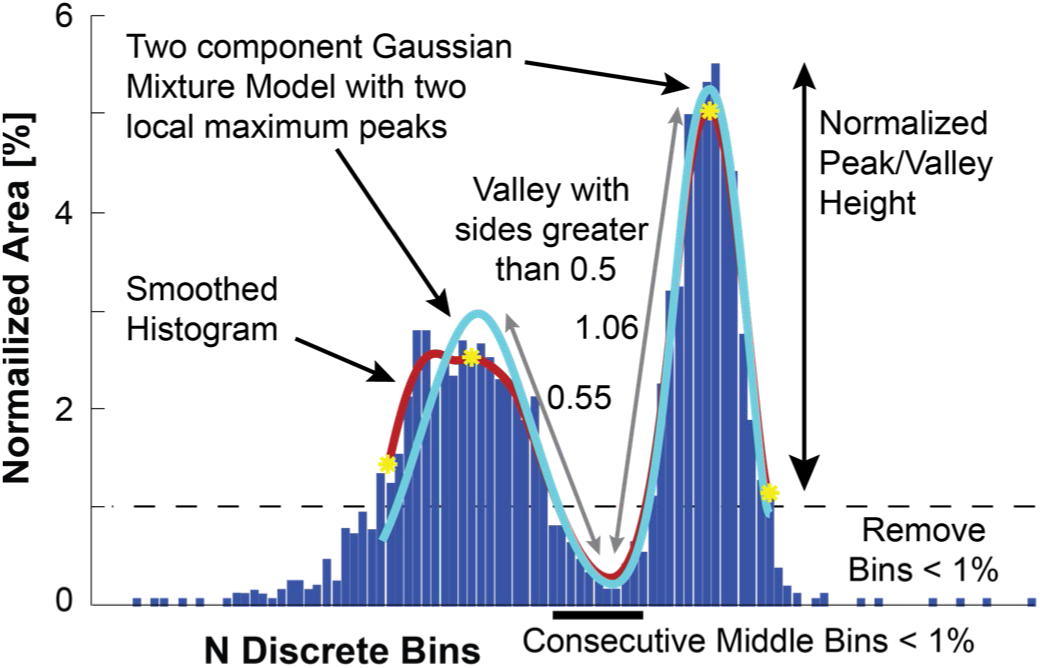
Multi-modality Detector. Generic function to determine if more than one mode (group) is present in a 1 dimensional data set. Histogram is formed from data in N bins (N is the square root of the number of data points (Mosteller. F. and Tukey (1977, p. 49)). Histogram is normalized by the sum of the bin area. A smoothed histogram generated by choosing each bin value as the median of the bin before and after it (Cocatre-Zilgien 1992); the output is filtered by a moving average (window size = 5 bins). All preceding and trailing bins less than 1% are removed from analysis. Three identifiers are used to find number of modes. First, a Gaussian Mixture Model is generated with M components (M is equal to the number of local max peaks of the smoothed histogram); second, if consecutive middle bins are found below the 1% threshold then two or more modes are present; third, a local valley is found with both sides greater than the normalized height of. 50 then two or more modes are present.

To indicate multi-modality within a set of data, the follow conditions were evaluated:

1. Any valley whose neighboring peaks are both greater than 0.50 is an indication the data is not unimodal.
2. For any valley whose neighboring peaks are both greater than 0.25, a Gaussian mixture model is generated using the number of local peaks as the number of components. Finding more than one local peak in the combined distribution is an indication the data is not unimodal.
3. Finding three or more consecutive middle bins which are less than the 1% histogram area is an indication these data are not unimodal.

## GRANTS

This work was funded by NIH Grant R01NS-72343 and DARPA cooperative agreement N66001-11-C-4171. Any opinions, findings and conclusions or recommendations expressed in this material are those of the author(s) and do not necessarily reflect the views of the Defense Advanced Research Projects Agency (DARPA) and SPAWAR System Center Pacific (SSC Pacific).

## DISCLOSURES

No conflicts of interest, financial or otherwise, are declared by the authors.

## AUTHOR CONTRIBUTIONS

D.B. analyzed data and designed algorithm and metrics; D.B., L.F., R.G., and D.W. interpreted results of experiments; D.B. prepared figures and drafted manuscript; D.B., L.F., R.G., and D.W. edited and revised manuscript; D.B., R.G. and D.W. approved final version of manuscript.

---

**Figure A2:**
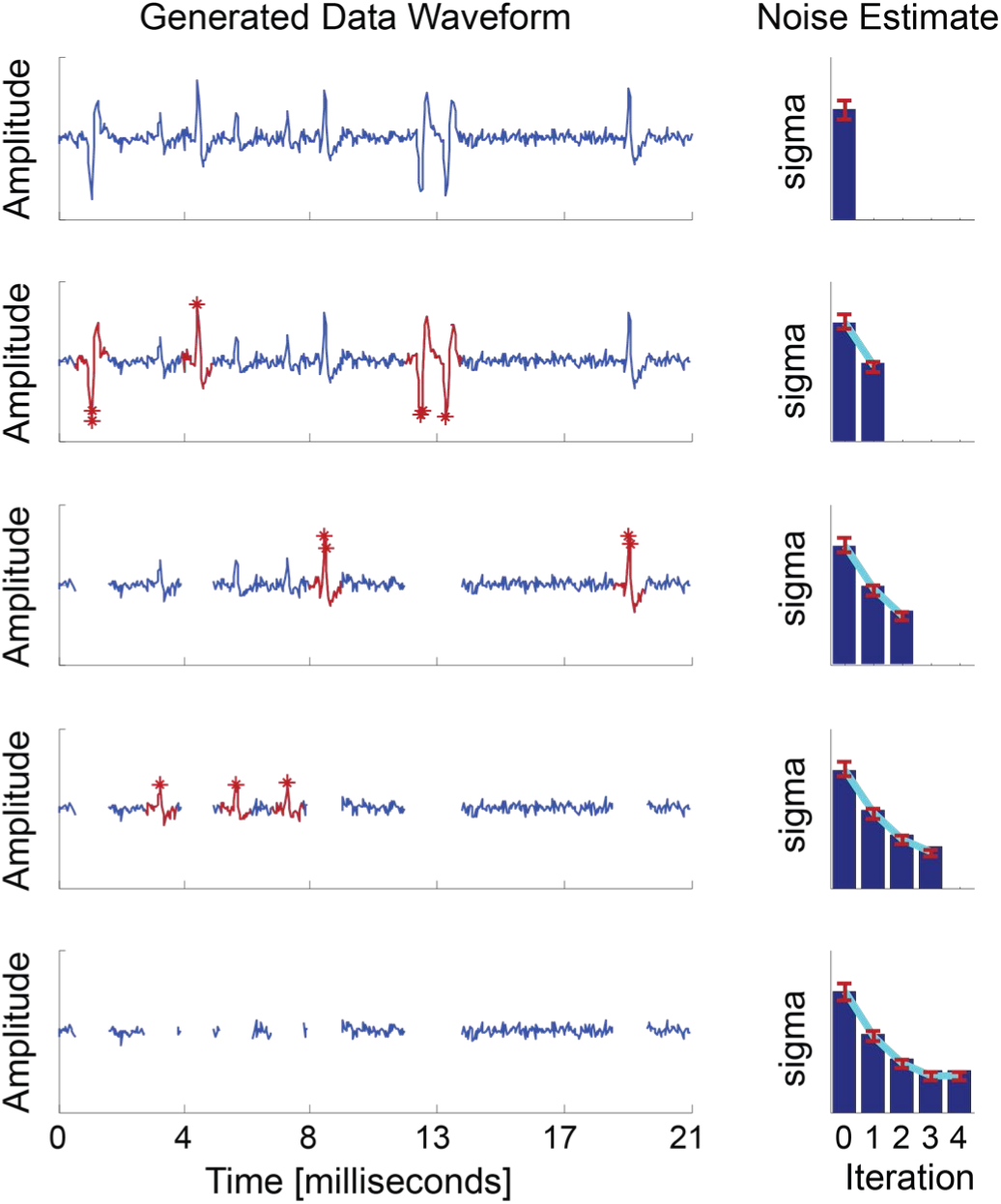
Noise estimation algorithm for determining SNR of neural signal. Iteration 0 shows the raw generated waveform. During each iteration, the standard deviation is estimated. The red dots indicate data points which lay more than four standard deviations from the mean of the entire waveform (shown in blue). A one millisecond snippet of the waveform is removed about that point (shown in red), and the next iteration estimates the updated distribution and removes more snips. This continues until the estimate for the standard deviation of the noise converges (shown on right column).

**Figure A3:**
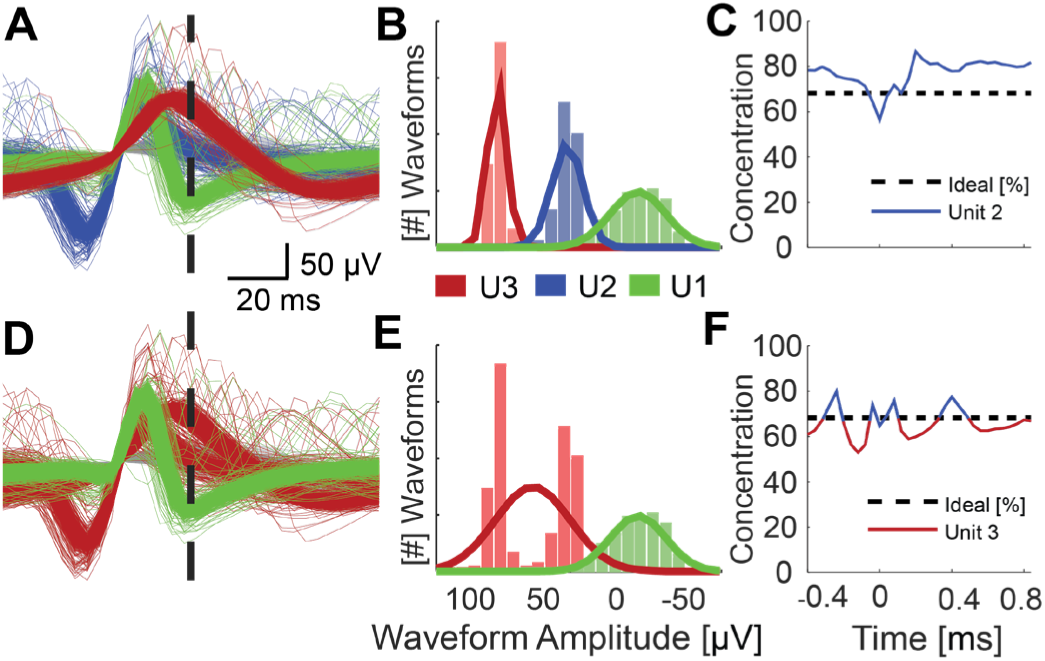
Residual Metric for Determining Over-sorted Units. Simulated data from the WaveClus dataset was used to generate the figures. (A, D) Histogram slices showing the distribution of waveform amplitudes (color-coded by Unit ID number) at various time stamps. (B) Histogram of waveform amplitudes (color-coded by Unit ID number) at time 0.1ms after threshold crossing. (C) Empirical measure of variability of the Gaussian fit (individually estimated at each sample point) for Unit 2. Concentration about the mean is the percent of waveforms within one estimated standard deviation. A lower empirical measure means greater variably than the estimated fit. Higher concentration about the average means a higher empirical measure. We see that nearly all sample points are empirically measured to have a lower variance (greater percent of waveforms) than the estimated fit (Ideal Sigma = 1). No sample point is multi-modal. This unit would pass the metric. (E) Histogram of an under-sorted Unit 3. Unit 3 contains information from both Neuron 1 and Neuron 2 from (B). We see the Gaussian curve poorly fits the histogram. (F) If any sample point with a greater variance (lower percent of waveforms) than the ideal is multi-modal, the unit fails the metrics. We see several sample points which are multi-modal and have a greater variance than the estimated fit. This unit would fail the metric.

**Figure A4:**
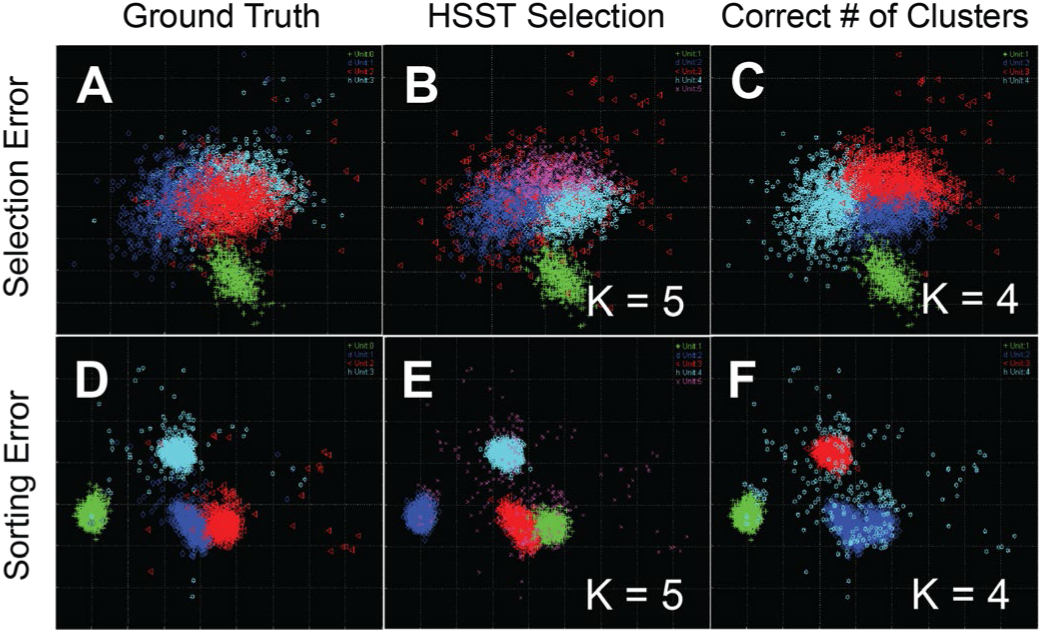
Visualization of Parameter Selection and Poor Sorting Errors. Waveforms are shown in 2D PCA space. Color denotes different clusters. Ground truth classification of waveforms shown (A) in different colors. 2D PCA space is used purely for visualization of data. The light blue line in Figure 6A shows the classification error rate as a function of the parameter selection. We see HSST selects ‘K=5’ as the optimal parameter value, even though ‘K=4’, yields the lowest classification error rate (B). Here we show (C) the classification of waveforms of the same light blue line from Figure 6A, this time at the parameter value “K=4”. Ground truth classification of waveforms shown in Figure 6B in different colors. Looking at the dark blue line from Figure 6B, we see HSST selects ‘K=5’ as the optimal parameter value (E), even though the number of clusters in the known dataset is 4. A parameter value of 5 yields the lowest classification error rate, and HSST selects it appropriately. Here we show the classification of waveforms of the dark blue light in Figure 6B at “K=5”. In Panel F, we show the classification of waveforms of the dark blue light in Figure 6B at “K=4”. Even though this is the “correct” number of clusters, the algorithm performance is poor.

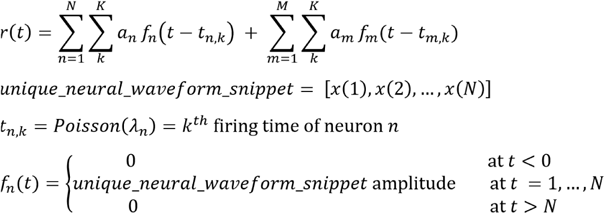

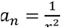 where *r*_*n*_ isthe distance from neuron *n* to the simukated electrode

Equation A1: NeuroCube Model. r(t) is the recorded voltage of the simulated electrode. 500 different unique neural snippets are available. Each neuron n, is randomly assigned one of the 500 unique waveforms. N primary neurons are specified by the user (N < 6). Each primary neuron is guaranteed to have a different unique waveform. M background neurons are auto generated at a distance greater than 250um with a density specified by the user (M < 10E+7).

**Table A1:** List of all fixed parameters in the HSST algorithm. All these were fixed for all analysis throughout the paper.

| Parameter | Location |
| --- | --- |
| Amount of temporal overlap in D'Prime distributions | Over-sorting Metric: Dissimilar Peaks |
| Amount of overlap in SSE curves | Noise Metric: ISI Exp. Fit |
| Threshold for signal data points | Noise Metric: SNR |
| Number of consecutive bins | Mutli-Modality Detector |
| Size of valleys | Multi-Modality Detector |

**Figure A5:**
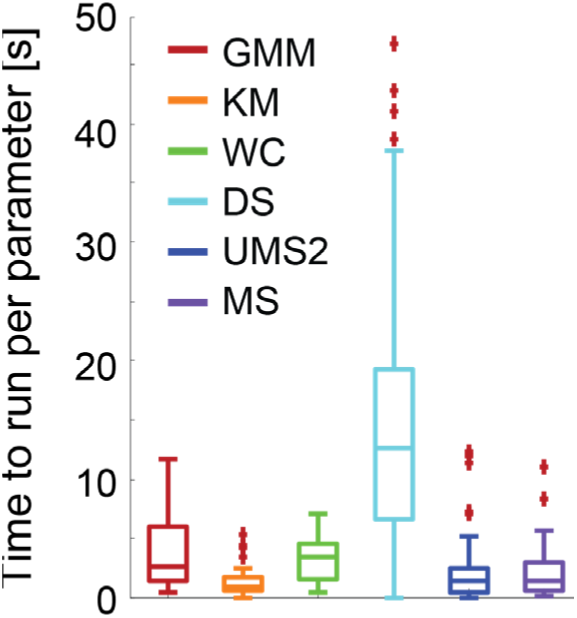
Algorithm Speed. Box plot of speed of each algorithm including the overhead of running HSST at the end of each parameter set (as you can see with MATLAB-KM method, the total time averages less than two seconds, so the additional time of scoring a sort with HSST is of negligible consequence). The Y Axis shows seconds to sort and score (using a single set of parameters) a dataset running the given algorithm. The 80 datasets sorted and

## Notes

https://github.com/davidbjanes/HSST

### Competing Interest Statement

The authors have declared no competing interest.

